# Listening difficulties in children: Behavior and brain activation produced by dichotic listening of CV syllables

**DOI:** 10.1101/721209

**Authors:** David R. Moore, Kenneth Hugdahl, Hannah J. Stewart, Jennifer Vannest, Audrey J. Perdew, Nicholette T. Sloat, Erin Cash, Lisa L Hunter

## Abstract

Listening difficulties (LiD), also known as auditory processing disorders, are common in children with and without hearing loss. Impaired interactions between the two ears have been proposed as an important component of LiD. Previous studies have focused on testing using multiple sequential dichotic digits that carry a substantial memory load and executive control demands. We examined the ability of 6-13 year old children with normal audiometric thresholds to identify and selectively attend to dichotically presented CV syllables using the Bergen Dichotic Listening Test (BDLT; www.dichoticlistening.com). Children were recruited as typically developing (TD; n=39) or having LiD (n=35) based primarily on composite score of the ECLiPS caregiver report. Different single syllables (ba, da, ga, pa, ta, ka) were presented simultaneously to each ear (6×36 trials). Children reported the syllable heard most clearly (non-forced, NF) or the syllable presented to the right (forced, FR) or left (FL) ear. Interaural level differences (ILDs) manipulated bottom-up perceptual salience. Dichotic listening data (correct responses, Laterality Index) were analyzed initially by group (LiD, TD), age, report method (NF, FR, FL) and ILD (0, ± 15 dB) and compared with speech-in-noise thresholds (LiSN-S) and cognitive performance (NIH Toolbox). fMRI measured brain activation produced by a receptive speech task that segregated speech, phonetic and intelligibility components. Some activated areas (planum temporale, inferior frontal gyrus and orbitofrontal cortex) were correlated with dichotic results in TD children only.

Neither group, age nor report method affected the Laterality Index of right/left recall. However, a significant interaction was found between ear, group and ILD. Children with LiD were more influenced by large ILDs, especially favoring the left ear, than were TD children. Neural activity associated with Speech, Phonetic and Intelligibility sentence cues did not significantly differ between groups. Significant correlations between brain activity level and BDLT were found in several frontal and temporal locations for the TD but not for the LiD group.

Children with LiD were more influenced by large ILDs, especially favoring the left ear, than were TD children and were thus less able to modulate performance through attention, and more driven by the physical properties of the acoustic stimuli.

## INTRODUCTION

Listening is often considered to be the active counterpart of passive hearing; “paying thoughtful attention to sound” (Keith et al., 2019); after Merriam-Webster). By definition, therefore, children with listening difficulties (LiD) may have problems with thought, attention or hearing. In practice, a considerable number of children seen at audiology clinics who have LiD are, on further testing, found to have normal audiograms, the pure-tone detection, gold-standard measure of hearing (Hind et al., 2011). For these children, a wide variety of symptoms are reported by caregivers (American Academy of Audiology, 2010; Moore & Hunter, 2013) that may be summarized as difficulty responding to meaningful sounds while ignoring irrelevant sounds. For at least 40 years, children with these symptoms have, following further testing, been diagnosed by some audiologists as having an auditory processing disorder (APD), but that diagnosis has not gained universal acceptance (Moore, 2018), so we will generally refer to the symptoms here by the more generic term of LiD.

Impaired interactions between the two ears have been proposed as an important component of LiD, based mainly on studies of dichotic listening (DL), the simultaneous presentation of different acoustic signals to the two ears (Broadbent, 1956; Keith, 2009; Kimura, 1961). However, other aspects of binaural interaction, including sound localization (Blauert, 1997) and binaural (Moore etn al., 1991; Pillsbury, Grose, & Hall, 1991) or spatial (Cameron & Dillon, 2007) release from masking have received substantial attention as contributors to LiD in adults and in children. Many other aspects of hearing and listening have also been studied in children with LiD (de Wit et al., 2016; Moore et al., 2010; Weihing et al., 2015; Wilson, 2018) leading, overall, to the emergence of two dominant hypotheses concerning the nature of the problem experienced by these children. The first, and more traditional hypothesis is that a disorder, central (C)APD, is primarily a result of impaired processing of auditory neural signals in the central auditory system, defined as the brain pathway from the auditory nerve to the auditory cortex (Rees & Palmer, 2010). The second, disruptive hypothesis is that LiD are due primarily to impaired speech/language synthesis, inattention, or other executive function impairment in cortical processing of auditory information beyond the central auditory system. The study reported here was motivated by an attempt to distinguish between these hypotheses.

Previous studies of DL in children have focused on listener reports of multiple sequential dichotic digits (Musiek, 1983) that carry a substantial memory and executive control load in addition to their linguistic and acoustic demands. Nevertheless, dichotic digits, often described confusingly as a test of binaural integration (American Academy of Audiology, 2010; Brenneman et al., 2017), has become one of the most common clinical tests of APD (Emanuel et al., 2011). Various dichotic digit-based training programs have been proposed as interventions for the remediation of APD (Moncrieff et al., 2017). However, recent research (Cameron et al., 2016) has questioned whether dichotic digits testing involves any binaural interaction. They found that performance on a *diotic* version of the test (presenting the same digits simultaneously to the two ears) correlates highly (r = 0.8) with performance on the dichotic version. The results suggest that, while binaural hearing may be disrupted during listening to dichotic digits, multiple, diverse abilities (acoustic discrimination, semantic identification, attentive listening, separation of two simultaneously presented sounds, accurate recall of heard digits) determine performance on these tasks.

As part of a larger Cincinnati Children’s Hospital program to investigate the nature of LiD in 6-13 year old (y.o.) children with normal audiometric thresholds (SICLID – Sensitive Indicators of Childhood Listening Difficulties), we examined those children’s ability to identify dichotically presented consonant-vowel (CV) syllables using the Bergen Dichotic Listening Test (BDLT; see www.dichoticlistening.com). In the BDLT (Hugdahl et al., 2009), two different CV syllables (from ba, da, ga, ka, pa, ta) are presented simultaneously, one to each ear, and the listener is asked either to report the first or most clearly heard syllable (non-forced, NF, condition), or selectively to report only that syllable presented to the left (forced left, FL) or to the right (forced right, FR) ear. In the NF condition, also known as the ‘Listen’ mode, the proportion of syllables presented to the right ear that is correctly reported consistently exceeds the proportion presented to the left ear that is correctly reported, a right ear advantage (REA). The REA is a robust, bottom-up, stimulus-driven, perceptual effect that is reflected in left-hemifield dominant activation of the auditory cortex (Hugdahl et al., 1999; Rimol et al., 2005; Westerhausen & Hugdahl, 2010). It is modulated by top-down, cognitive influences (attention, executive function, working memory, training), reflected in FR and, particularly, FL performance. An acoustic interaural level difference between the syllables can offset the REA and thus serve as a physical measure (in dB) of a cognitive construct (Hugdahl et al., 2008; Westerhausen et al., 2009). For these reasons, as well as its simplicity and the extensive literature on it, the BDLT is well-suited to investigate neural processes of listening in children. This study represents the first, to our knowledge, where BDLT has been used to examine children with LiD/APD.

Previous studies have shown that BDLT ‘Concentrate’ modes (FL, FR) activate different brain regions in adults when contrasted with the Listen mode (Westerhausen & Hugdahl, 2010). Thus, FR activates a ‘dorsal attention network’, consisting of the rDLPFC and, weakly, lDLPFC and the bilateral occipital cortex. FL activates a ‘cognitive control network’, consisting of the bilateral angular gyrus, DLPFC and anterior cingulate cortex (Westerhausen et al., 2010). We have taken another approach to examine brain activation produced by the BDLT, using a sentence listening and speaker identification test to produce BOLD activation inside a 3T MRI scanner. Here, we correlated the sentence-induced brain activation in a way that dissects out components of receptive speech (speech, phonetics, intelligibility). We examined activation for each group (TD, LiD) of children’s performance by BDLT listening mode (NF, FL, FR) and interaural acoustic bias (interaural level difference, ILD).

Testing the hypothesis that children with LiD have problems with cortical language, attention and executive function beyond the central auditory system, the predictions of this study were that (i) children with LiD will perform normally on BDLT in Listen mode but will have difficulty in Concentrate mode, based on their overall tendency to perform poorly on cognitive tasks despite normal hearing; and (ii) children with LiD will have atypical brain activation associated with top-down processing of speech (intelligibility, speech), but not non-speech sounds (phonetics, acoustics) correlated with BDLT performance. To investigate these predictions, we examined BDLT performance in normally hearing 6-13 y.o. children and correlated that performance with other tests of speech perception, cognition, and speech sound-evoked fMRI.

## MATERIALS AND METHODS

### Participants

Children with LiD were recruited initially from a medical record review study of over 1,100 children assessed for APD at Cincinnati Children’s Hospital (CCH; Moore et al., 2018). Caregivers of children diagnosed with APD (including those with a “Weakness”) who responded to invitation to participate were sent questionnaires including the ECLiPS, below, and a background questionnaire. Those who completed and returned the questionnaires were invited to bring their child into the lab for a study visit. Over time, recruitment expanded to include the use of CCH IRB-approved materials, advertising, and messages via print, electronic, social and digital media at hospital locations and in the local and regional area for participation of families with children who had a “listening difficulty,” or were “without any known or diagnosed learning problem”. Following a positive response and a brief telephone interview to screen for listening status, families were sent the same questionnaire pack and were invited for a study visit as described below.

Seventy four children aged 6-13 y.o. completed BDLT testing and secondary behavioral testing. All of these children had normal hearing, bilaterally, defined as clear ears, A-type tympanometry, and pure tone thresholds ≤20 dB at octave frequencies between 0.25 – 8 kHz (Figure 1A) using standard audiometric procedures. Additional, extended high frequency audiometry (10 – 16 kHz; Figure 1B) was also obtained, but inclusion did not require any criterion level of performance at these frequencies. Seventy children received MRI scanning (95%).

**FIGURE 1:**
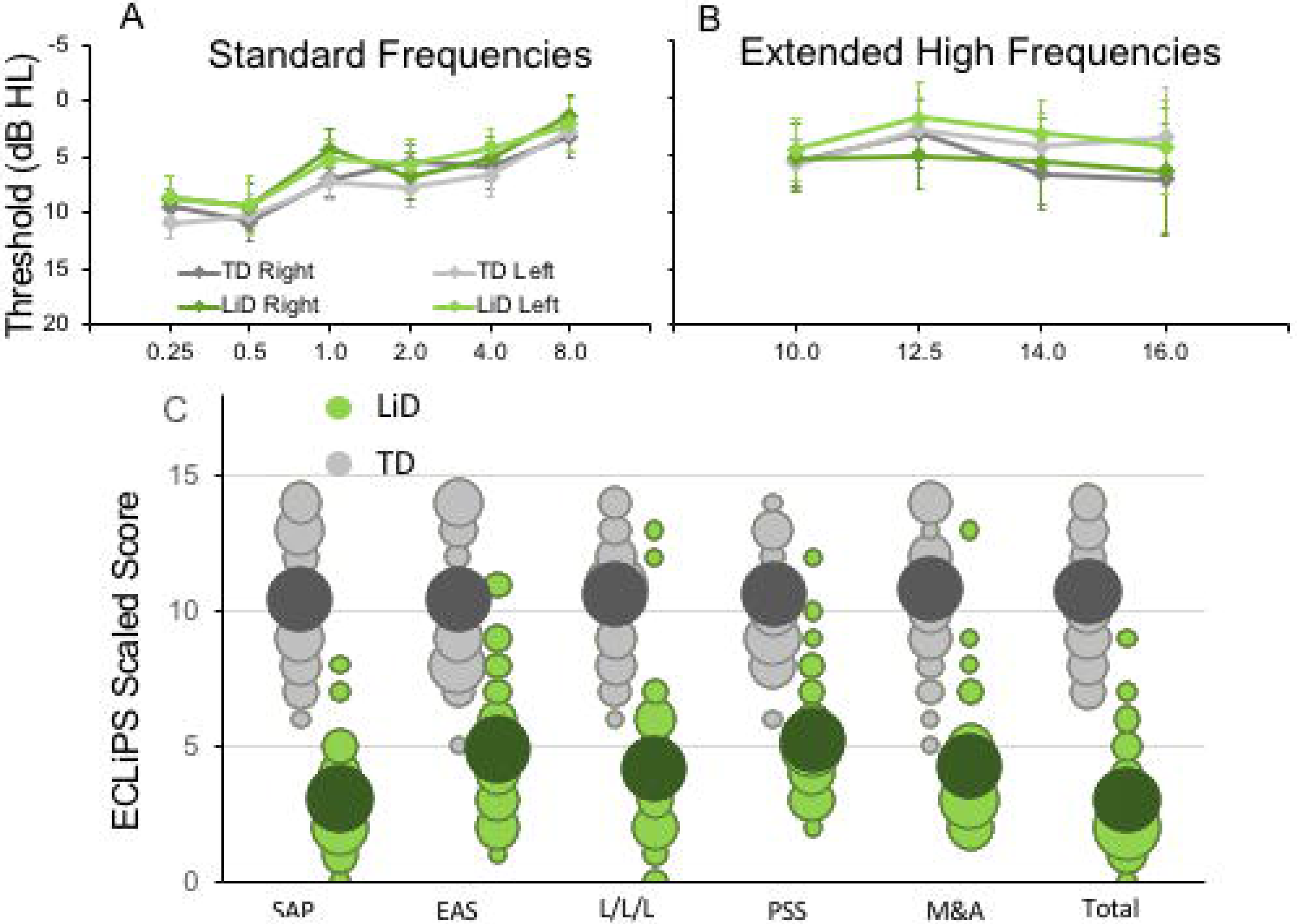
Children in study had no hearing loss but reduced caregiver evaluation of listening skills. **(A)** Mean (± 95% CI) pure tone audiometric thresholds at standard frequencies (0.25 – 8.0 kHz) for children in each group. TD – Typically developing, LiD – Listening difficulties. **(B)** Audiometric thresholds at extended high frequencies. **(C)** Caregiver evaluation using the ECLiPS (Barry & Moore, 2015; Barry et al., 2015). Scaled scores (normalized to mean = 10, s.d. = 3) shown separately for Total score, Speech & Auditory Processing (SAP), Environmental & Auditory Sensitivity (EAS), Language/Literacy/Laterality (L/L/L), Memory & Attention (M&A), Pragmatic & Social Skills (PSS). Bubble size proportional to number of children achieving each scaled score.

The ECLiPS questionnaire (Barry & Moore, 2015; Roebuck & Barry, 2018) is a 38 item inventory asking users to agree or disagree (5 point Likert scale) with simple statements about their child’s listening and related skills. Total standardized ECLiPS scores ≥5 designated typically developing (TD), and scores <5, or a previous diagnosis of APD, designated LiD, resulting in 39 TD children and 35 children with LiD (Figure 1C).

Demographics, audiological status, secondary testing of auditory and speech perception, and cognitive performance of the larger SICLID sample (n = 146) will be reported in greater detail elsewhere.

### Behavioral tests

#### Bergen Dichotic Listening Test (BDLT)

Digitally recorded test materials were provided by the Department of Biological and Medical Psychology, University of Bergen, Norway. Test materials and general procedures are described in detail elsewhere (Hugdahl et al., 2009). Listeners were seated in a sound treated booth and instructed to attend to and verbally repeat speech sounds presented via Sennheiser HD 25-1 headphones connected to a laptop PC. Control software was Direct RT. Two different CV syllables from a list of six (/ba/, /da/, /ga/, /ka/, /pa/, /ta/) were presented simultaneously, one to each ear at a level of 65 dB SPL. Each trial was started manually by the tester when the participant was ready. In a ‘non-forced’ (NF) condition the listener was asked to report the syllable they “heard best”. Alternately, the listener was asked to report only that syllable presented to the left (forced left, FL) or to the right (forced right, FR) ear.

A test session started with 12 practice trials (NF). For the first 6 of these (ILD = 0 dB), the listener had to repeat one of the syllables correctly in 5/6 trials to proceed. For the second 6 trials, ILD varied between +15 (right louder) through −15 to 0 dB each two trials, and the listener had again to get 5/6 trials correct to proceed. The practice trials were repeated if a listener did not achieve the prescribed correct response rate. Five children (in addition to the 74) were excused from the experiment when they failed to achieve the prescribed correct response rate after several runs. Data collection sessions (x6) each consisted of 36 trials containing every possible pair combination. The first two, NF sessions had 12 trials each of ILD = 0, +15, −15. In randomized order, there followed two FR and two FL sessions, with 12 trials each of ILD = 0, −15, +15 dB (first session), and ILD = 0, +15, −15 dB (second session). A short break was provided between each session. Data were downloaded to REDCap (Harris et al., 2019; Harris et al., 2009) for storage and analysis.

#### LiSN-S

The Listening in Spatialized Noise – Sentences (LiSN-S) task (Cameron & Dillon, 2007) (www.LiSN-S.com) measures ability to attend, hear and recall sentences in the presence of distracting sentences. LiSN-S was administered using a laptop, a task-specific soundcard, and Sennheiser HD 215 headphones. Participants were asked to repeat a series of target sentences (‘T’), presented directly in front (0°), while ignoring two distracting talkers. There were four listening conditions, in which the distractors change voice (different or same as target) and/or (virtual) position (0° and 90° degrees relative to the listener). The test was adaptive; the level of the target speaker decreased or increased in signal/noise ratio (SNR) relative to the distracting talkers as the listener responded correctly or incorrectly. Testing continued for a minimum of 22 trials per condition (including 5 practice items that did not contribute to the score). Testing stopped when SEM < 1 or after 30 trials. The 50% correct SNR was either the ‘Low cue speech reception threshold’ (SRT; same voice, 0° relative to the listener) or the ‘High cue SRT’ (different voice, 90° degrees relative to the listener). Three ‘derived scores’ were the Talker Advantage, Spatial Advantage, and Total Advantage, so-called because each is the difference between SRTs from two conditions.

#### NIH Cognition Toolbox

Cognition was assessed using the NIH Toolbox - Cognition Domain battery of tests (Weintraub et al., 2013). Participants completed testing online or via iPad app in accordance with the current NIH recommendations in a private sound attenuated booth or quiet room. The battery contains up to seven different standardized cognitive instruments measuring different aspects of vocabulary, memory, attention, executive functioning, etc. The precise composition of the testing battery is dependent on participant age. Sixty five participants in this study completed the picture vocabulary test (TPVT), flanker inhibitory control and attention task (Flanker), dimensional change card sort test (DCCS), and picture sequence memory test (PSMT). Each test produced an age-corrected standardized score and the scores of all four tests were combined to calculate a single, Early Childhood Composite. Additional tests, contributing to the Crystallized, Fluid and Total Composite scores, were the List Sorting Working Memory test (LSWM), the Pattern Comparison Processing Speed test (PCPS), and the Oral Reading Recognition test (RR).

TPVT is an adaptive test in which the participant is presented with an audio recording of a word and selects which of four pictures most closely matches the meaning of the word. In the Flanker, testing inhibition/attention, the participant reports over 40 trials the direction of a central visual stimulus (left or right, fish or arrow) in a string of five similar, flanking stimuli that may be congruent (same direction as target) or incongruent (opposite direction). The DCCS tests cognitive flexibility (switching attention). Target and test ‘card’ stimuli vary along two dimensions, shape and color. Participants are asked to match test cards to the target card according to a specified dimension that varies for each trial. Both the Flanker and DCCS score accuracy and reaction time. PSMT assesses episodic memory by presenting an increasing number of illustrated objects and activities, each with a corresponding audio-recorded descriptive phrase. Picture sequences vary in length from 6-18 pictures depending on age, and participants are scored on the cumulative number of adjacent pairs remembered correctly over two learning trials.

### Magnetic Resonance Imaging

#### Stimuli and task

fMRI scanning included an active speech categorization task. Sixteen BKB sentences (Bench et al., 1979) recorded by a single male North American speaker under studio recording conditions were presented using sparse scanning procedures (‘HUSH’; (Deshpande et al., 2016; Schmithorst & Holland, 2004). Specifically, sentences were presented during a 6-second silent interval followed by 6 seconds of fMRI scanning (details below). Following methods described by Scott and colleagues (Scott et al., 2000), recordings limited to < 3.8 kHz but otherwise unprocessed were delivered as ‘Clear’ speech sentences. ‘Rotated’ speech stimuli were created by rotating each sentence spectrally around 2 kHz using the (Blesser, 1972) technique. Rotated speech was not intelligible, though some phonetic features and some of the original intonation were preserved. ‘Rotated and Vocoded’ speech stimuli were created by applying 6 band noise-vocoding (Shannon et al., 1995) to the rotated speech stimuli. While the rotated noise-vocoded speech was completely unintelligible, the character of the envelope and some spectral detail was preserved. The listener’s task was to make a button press after each sentence presentation, indicating whether a cartoon image (‘human’ or ‘alien’) matched the speaker of the sentence. In familiarization trials, before scanning, the clear speech was introduced as ‘human’ and the rotated/vocoded speech as ‘alien’. Each participant completed 3 practice trials with verbal feedback from the tester. If a trial was completed incorrectly, the stimuli and instructions were reintroduced until the listener showed understanding.

#### Procedure

All listeners wore foam ear plugs to attenuate the scanner noise, but they were still able to hear clearly the stimuli delivered binaurally (diotically) via MR-compatible circumaural headphones. Listeners completed 48 matching trials, 16 of each sentence type, with no feedback. To maintain scanner timings, the sentence task continued regardless of whether a response was made. However, if a response was not made on three trials in a row, the tester provided reminders/encouragement over the scanner intercom between stimuli presentations.

#### Imaging

MRI was performed using a 3 T Phillips Ingenia scanner with a 64-channel head coil and Avotec audiovisual system. The scanning protocol included a T1-weighted anatomical scan (1mm isotropic resolution) and the fMRI task described above using a sparse acquisition approach (‘HUSH’; TE = 30 ms, TR = 2000 ms, voxel size = 2.5 × 2.5 × 3.5 mm, 39 slices).

### Analysis

#### Behavioral Analysis

ECLiPS, LiSN-S and NIH Toolbox data were separately analyzed in two-way mixed effects ANOVA, with the Group variable (TD/LiD) and within-subject variables for subtests. Separate t-tests were used to examine composite scores.

Dichotic listening data were first analyzed in a four-way mixed effects ANOVA, with the variables 2 Groups (TD/LiD) x 2 Ear (Right, Left) x 3 Attention (NF, FR, FL) x 3 ILD (0, +15, −15), and number of correct reports as the dependent variable. The Group variable was treated as a between-group variable, while the ear, intensity and attention variables were treated as within-subject variables. In a second three-way ANOVA, we reduced the design to the variables Group x Attention x Intensity, and with the laterality index score as dependent variable. The laterality index (LI) score controlled for differences in overall performance between the participants, and was calculated according to the formula [(REar – LEar) / (REar + LEar) x 100].

To elucidate differences between groups in sensitivity to manipulating the physical acoustic environment of the stimuli, a third, two-way ANOVA further reduced the variables to Group x ILD, again based on the LI scores. Follow-up post hoc tests of main- and interaction-effects were done with Fisher’s LSD test. Significance threshold was set at p = 0.05 for all tests.

Correlations between DL, care-giver report, spatialized listening and cognitive function were conducted using Pearson’s coefficient between age-corrected DL-LI across ILD, and ECLiPS Total Score, LiSN-S Low Cue and Talker Advantage, and NIH Toolbox Total Composite.

#### Imaging analysis

First-level fMRI data were processed using FSL (FMRIB Software Library, https://fsl.fmrib.ox.ac.uk/fsl/). Anatomical T1 data and functional data were first reoriented using FSL’s *fslreorient2std*. Next, the T1 data were brain extracted using FSL’s BET. The brain extracted T1 image was then normalized and resampled to the 2mm isotropic MNI ICBM 152 non-linear 6th generation template using FSL’s FLIRT. For the functional data, the initial 3 time points were discarded to allow protons to reach T1 relaxation equilibrium. Slice timing correction and brain extraction were carried out using FSL’s “*slicetimer*” and BET, respectively. Outlying functional volumes were detected using FSL’s “*fsl_motion_outliers*” with the default RMS intensity difference. Cardiac and respiration signals were regressed out using AFNI’s “*3dretroicor*”. Motion correction of the BOLD time-series was carried out using MCFLIRT. Motion-related artifacts were regressed from the data by setting up a general linear model (GLM) using 6 motion parameters. The amount of motion during the scans did not differ between groups. The results were interpolated to a 2 mm isotropic voxel size and aligned to the Montreal Neurological Institute (MNI) template by first co-registering it with the participant’s T1 using FSL’s FLIRT.

Second level analysis was also conducted using FSL. A GLM approach was used to create group activation maps based on contrasts between conditions for all participants (i.e. regardless of LiD/TD status). Group composite images were thresholded using a family-wise error correction (p < 0.001) and clustering threshold of k = 4 voxels. Three BOLD activation contrasts were used to search for brain loci responding to different aspects of speech perception (modified from Scott et al., 2000). First, the ‘Speech’ activation map contrasts a signal having intelligibility, intonation, phonetics and sound with one lacking all these attributes except sound (clear > rotated/vocoded). Second, the ‘Intelligibility’ activation map contrasts a signal having all attributes with one retaining intonation, phonetics and sound (clear > rotated). Third, the ‘Phonetics’ activation map contrasts a signal having intonation, phonetics and sound with one having only sound (rotated > rotated + vocoded).

#### ROI Analysis

These three activation maps were used to identify brain regions showing significantly increased activation for speech, phonetic features and intelligibility. These active regions were used as ROIs for correlation analysis with the DL behavioral data within which significant group differences between TD and LiD were hypothesized. Statistical analysis used JASP (v. 0.10.2) to plot data regressions and calculate correlations.

## RESULTS

### Audiometry and caregiver report

No significant difference in pure tone auditory threshold detection was found between children who were TD and those who had LiD across either the standard (Figure 1A) or extended (Figure 1B) frequency range. Children formed a continuum of listening abilities, as assessed by caregivers, but two groups, TD and LiD were segregated, primarily on their total score on the ECLiPS (Figure 1C). Two children in the LiD group who overlapped with the TD range of scores, and an additional 11 children with LiD had a clinical diagnosis of APD.

### Auditory perceptual and cognitive function

Listening to sentences in ‘spatialized’ noise (LiSN-S) was significantly (p < 0.01) impaired in children with LiD on the Low Cue and High Cue conditions, and the derived Talker Advantage measure (Figure 2A). This pattern of results suggested that the children with LiD had problems with both the procedural demands and the specifically auditory demands of the task.

**FIGURE 2:**
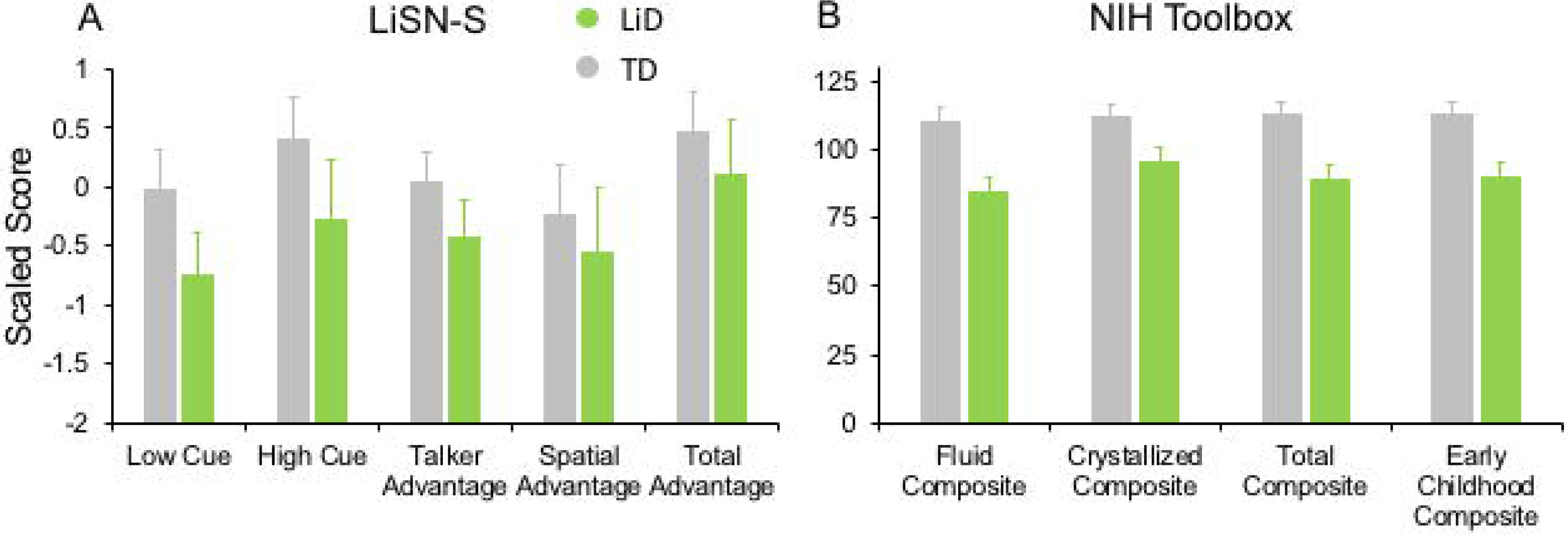
Children in LiD group had reduced speech in noise (LiSN-S) and cognitive (NIH Toolbox) performance relative to TD children. **(A)** LiSN-S (Cameron & Dillon, 2007) mean z-scores (± 95% CI) for each group. Low Cue (speech and distractors same direction, same talker), High Cue (speech and distractors different direction, different talker), Talker advantage (benefit re Low Cue of changing distracting talker), Spatial advantage (benefit re Low Cue of changing distractor position). Further details in text. **(B)** NIH Toolbox. Mean composite scores (normalized to mean = 100, s.d. = 15) for each group. Further details in text.

Related to the disability of children with LiD to perform the listening task (LiSN-S), we found they also had impaired performance on all subtests of the NIH Toolbox, summarized in Figure 2B (all p < 0.001). The mean standard score of the children with LiD was poorest on the Fluid Composite, composed of the visually-based NIH Toolbox DCCS, Flanker, PSM, LSWM and PCPS subtests. Performance was also significantly impaired on the Crystallized Composite, consisting of the PV and RR subtests. The PV was the only subtest on which success was partly dependent on auditory perception and receptive language function. However, results for the PV subtest (mean difference between LiD and TD groups = 15.9 points) were similar to those of the RR subtest (16.0 points). It therefore appears that the LiD group had a generalized, multi-modal mild cognitive impairment relative to the TD group.

### Dichotic listening

Children of all ages were generally able to complete the full 216 trials of BDLT testing in about 30 minutes, although there was a significant attrition rate as testing continued since the task is not the most engaging and fatigue was commonplace in both groups. Participants with LiD were more likely become frustrated or upset by the task. Frequent check-ins with the participant were needed and short breaks (a few minutes) were not uncommon. However, neither fatigue nor inattention were a basis for exclusion. Forced conditions were counterbalanced.

We first examined the BDLT results of all children in terms of number of syllables correctly identified, with a maximum possible score under each condition of 24 (12 trials x 2 blocks. 3 ILDs x 3 Attention conditions; Figure 3). For no ILD (ILD 0 dB), all three attention conditions (NF, FL, FR) showed a right ear advantage in both groups (Figure 3A). That advantage became larger for ILD +15 dB (Figure 3B) and reversed for ILD −15 dB (Figure 3C), all as expected from the DL literature, except that a REA was obtained even in the FL condition. For the ILD −15 dB condition, it appeared that the ear differences were smaller for the TD than for the LiD group. An overall four-way ANOVA was first run with the factor ‘Age’ as covariate to control for the small age difference between the groups (see below). This analysis showed a significant three-way interaction of ILD by ear by group: F(2, 142) = 5.70, p = 0.004, eta^2^ = .07. The interaction was followed-up with Tukey’s HSD test which showed that, while both groups were able to shift to a significant left ear advantage during the ILD −15 dB condition, this ability was exaggerated in the LiD group, controlling for multiple comparisons. The LiD group were thus less able to modify the acoustic ear advantage through cognitive influences.

**FIGURE 3:**
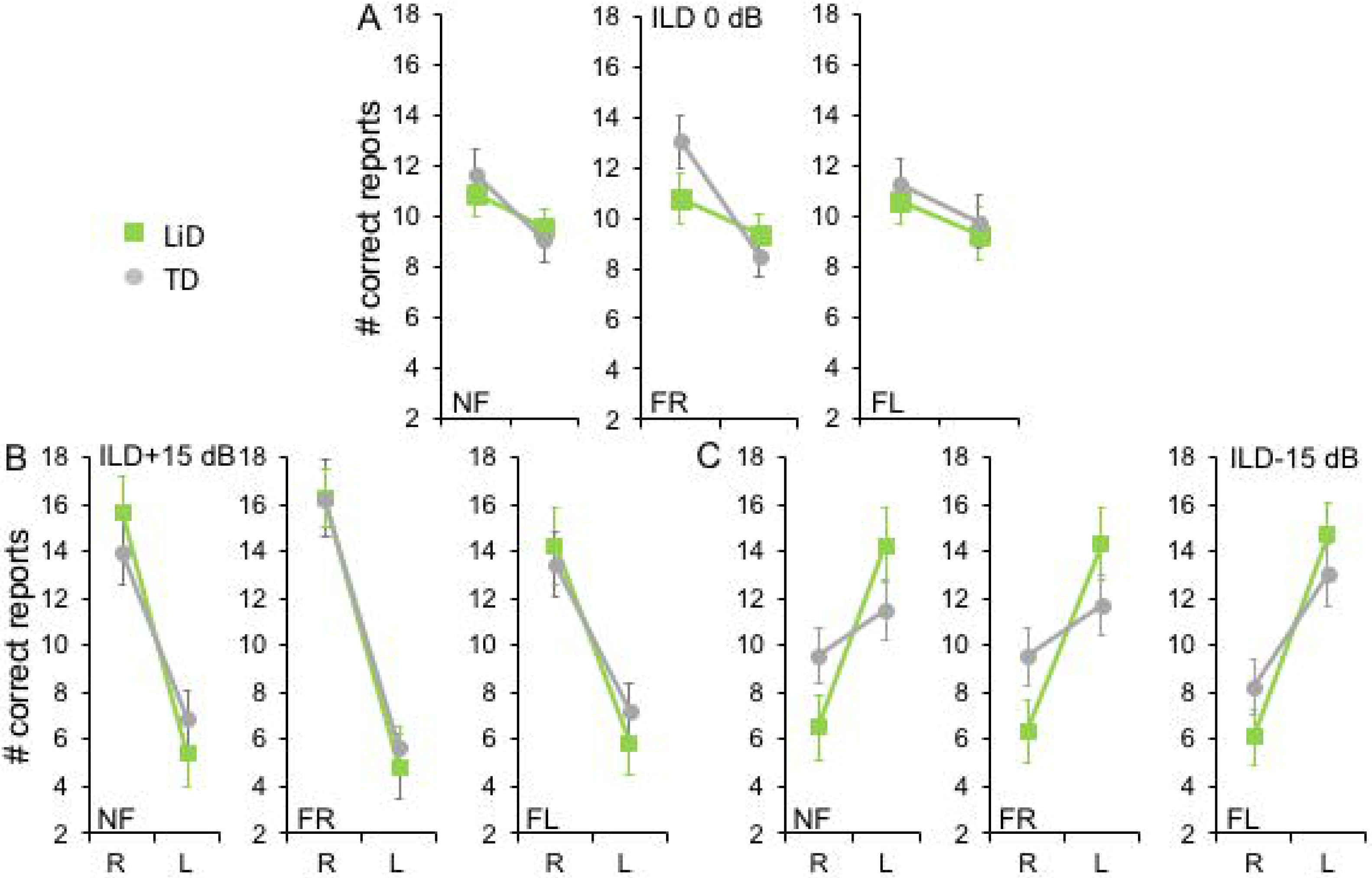
Children in both groups showed a right ear advantage in BDLT for correct syllable identification. Mean (± sem) number of correctly identified digits delivered to each ear as a function of attention condition (NF, FL, FR), group (LiD, TD), ILD, and ear (L, R). **(A)** ILD 0 dB. Same level of stimulus in each ear. **(B)** ILD +15 dB. Stimulus 15 dB more intense in right ear. **(C)** ILD −15 dB. Stimulus 15 dB more intense in left ear.

To investigate group differences further, we next examined the Laterality Index (LI; Figure 4A). Three-way ANOVA showed a significant effect of ILD: F(2,142) = 4.45, p = 0.013, partial eta^2^ = 0.06. There was a significant two-way interaction of ILD by group (Figure 4B): F(2,142) = 6.87, p = 0.001, partial eta^2^ = 0.08. Tukey’s HSD test showed significantly higher Laterality in the LiD group in the ILD −15 dB condition, controlling for multiple comparisons. Also shown on Figure 4B are typical young adult data (NF condition) from the study of (Westerhausen et al., 2009). At ILD 0 dB, the LI (REA) was smallest for the LiD group (6%), larger for the TD group (13%), and largest for the Adults (28%). Note that Westerhausen’s adult data were near parallel with the LiD data, but that the LiD data showed a stronger left ear influence at each ILD. Asymmetry of LI between the ILD ±15 dB was more marked for the TD than the LiD group, with TD children, like adults, showing a much larger LI for ILDs favoring the right ear. By contrast, children with LiD had larger but near symmetric LIs for ILD ±15 dB. Both groups of children showed different immature response patterns.

**FIGURE 4:**
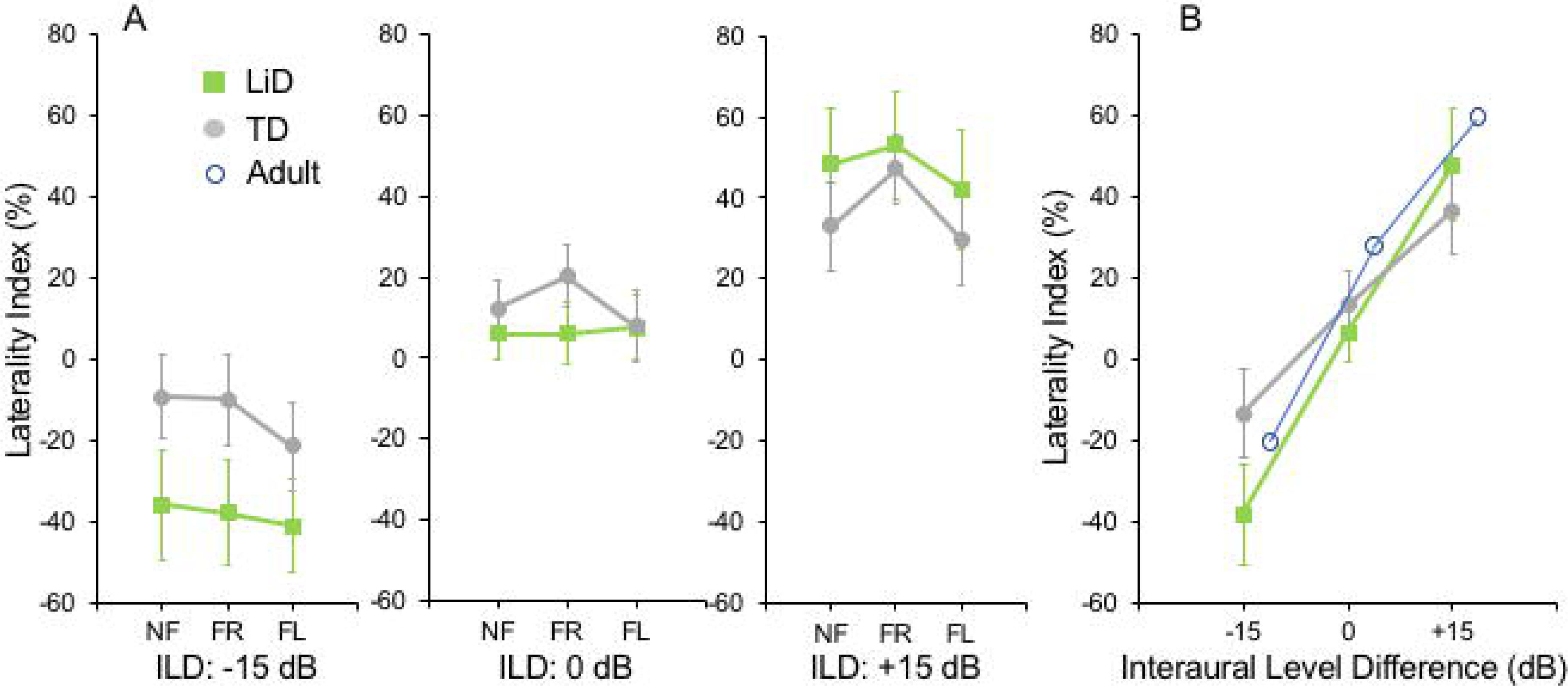
BDLT laterality index varied more in LiD group than in TD group as a function of ILD. **(A)** Same comparisons as in Fig. 3 expressed as mean (± 95% CI) percentage correct responses right ear relative to left ear (see text). **(B)** Mean (± 95% CI) LI as a function of ILD averaged across attention conditions in each group. Adult data from (Westerhausen et al., 2009).

LIs for three age groups, with both TD and LiD children together and divided approximately to equalize the number of children in each group, are shown in Figure 5. As above, small differences were seen between the age groups, with older children overall having slightly larger unsigned LIs than younger children, although not significant, as indicated from 3-way ANOVA with age x attention x ILD as variables. Note the positive LIs in the ILD 0 dB FL condition.

**FIGURE 5:**
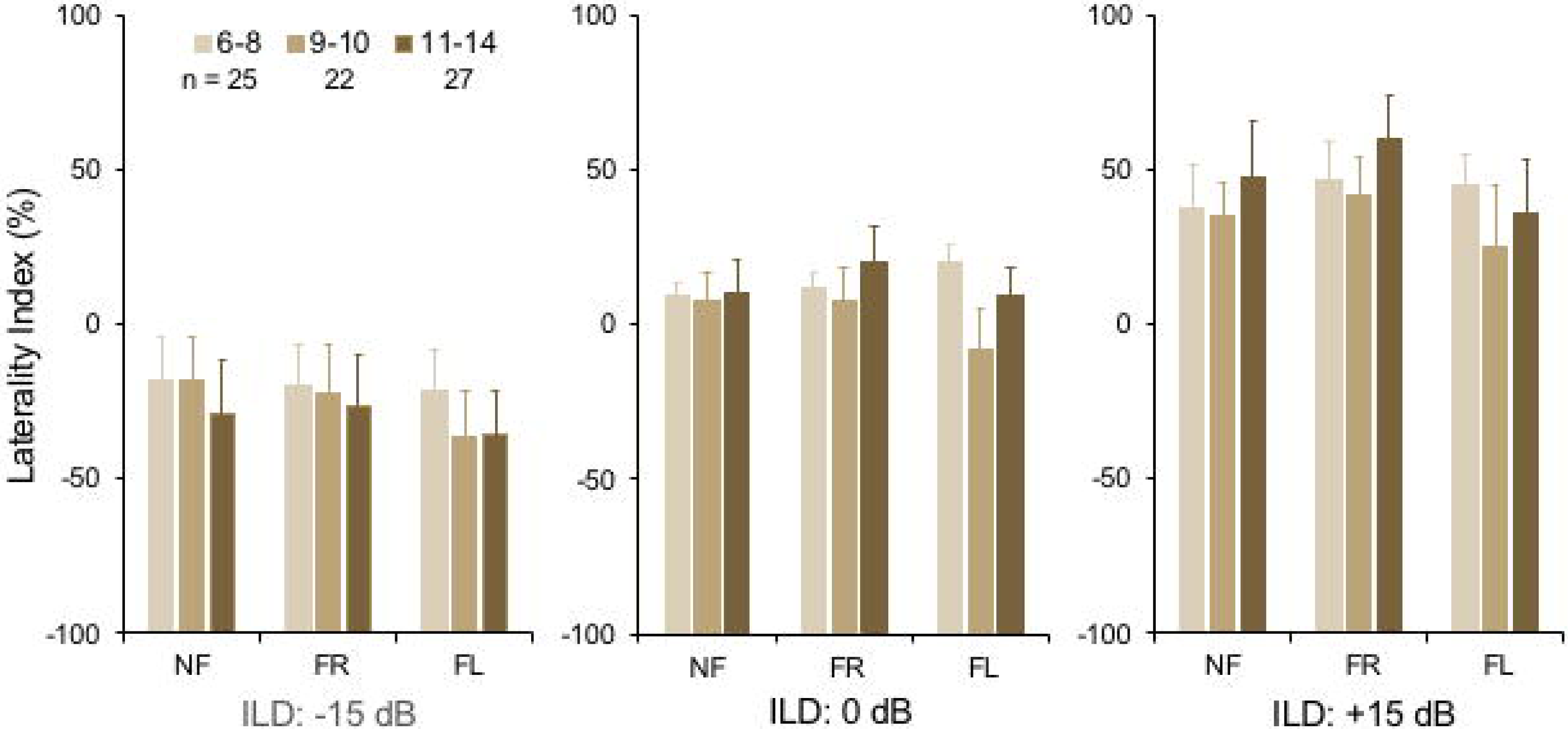
BDLT laterality index increased slightly, but non-significantly with age. Same comparisons as in Figs. 3 and 4; mean (± 95% CI) LI averaged across groups. Note that REA did not increase significantly across this age range.

In summary, neither group, age nor attention condition affected the Laterality Index of right/left recall. However, a significant interaction was found between group (LiD, TD) and interaural level difference (ILD). Children with LiD were more influenced by large ILDs, especially favoring the left ear, than were TD children and were thus less able to modulate performance through attention, and more driven by the physical properties of the acoustic stimuli.

### Correlations between behavioral measures

Few significant correlations were observed between the ECLiPS, the LiSN-S or the NIH Toolbox data, and DL-LI measures. From a total of 99 comparisons, only 6 LiSN-S and Toolbox measures were significant at p < 0.01 (Figure 6), uncorrected for multiple comparisons. For the ECLiPS (not shown), only 3/9 comparisons were significant at p < 0.05, and all 3 comparisons were for ILD −15 dB, at which LIs and differences between groups were largest (Figure 4B). Similar patterns were seen for the LiSN-S and the Toolbox (Figure 6). Toolbox data showed the strongest and most consistent correlations (Figure 6B). For example, the Fluid Composite was associated with LI (p < 0.001); all four Toolbox measures showed low cognitive performance associated with strongly negative LI, reaching in some mostly LiD cases −100%, a strong left ear advantage.

**FIGURE 6:**
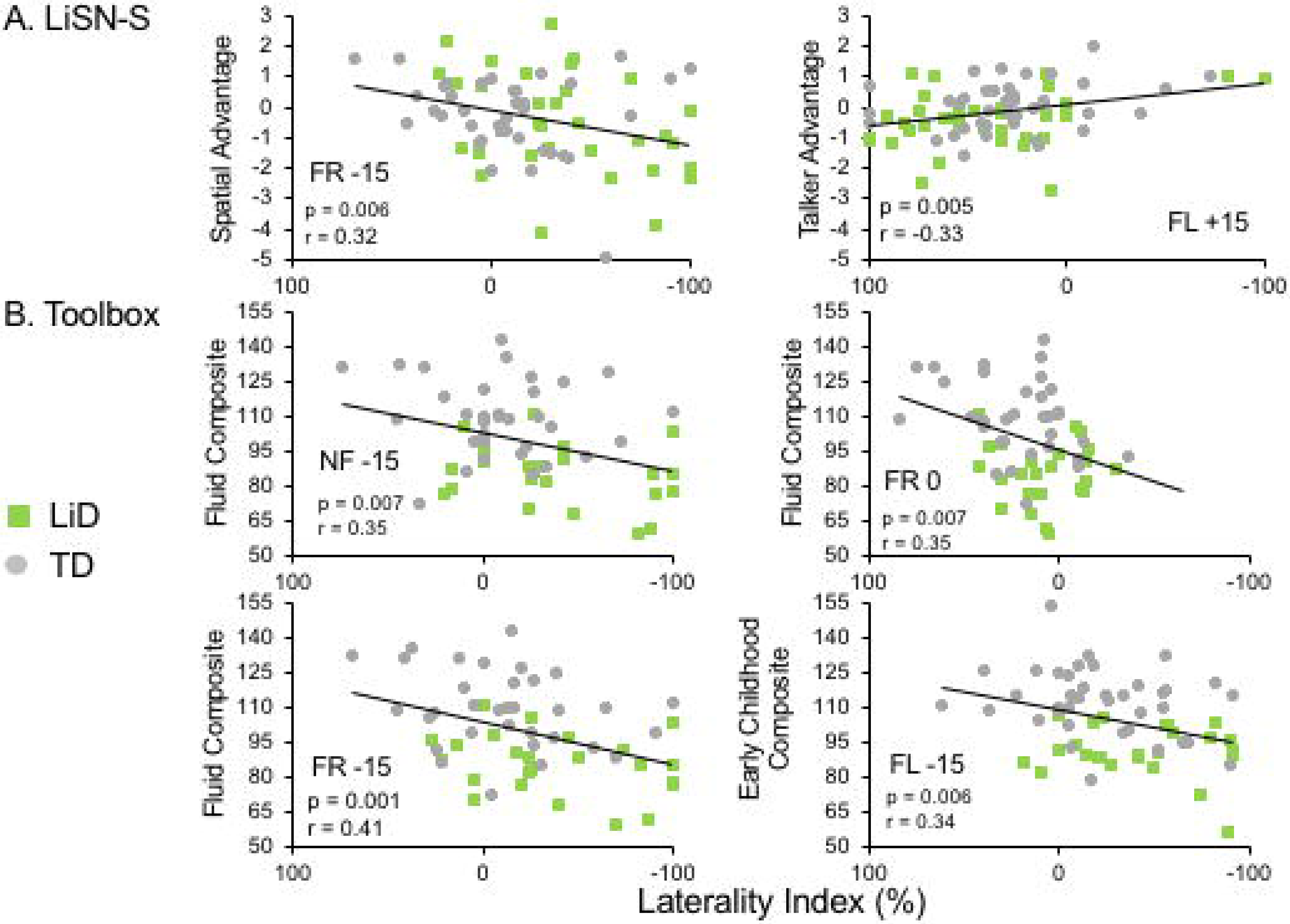
Limited correlations were seen between LI and behavioral hearing and cognitive tests. Comparative performance of individuals in both groups on **(A)** Lisn-S Advantage measures and (B) NIH Toolbox composite measures.

### fMRI

All children performed well in the scanner on the active speech categorization task, although the TD children performed more accurately, and with shorter reaction times, than those with LiD (Figure 7).

**FIGURE 7:**
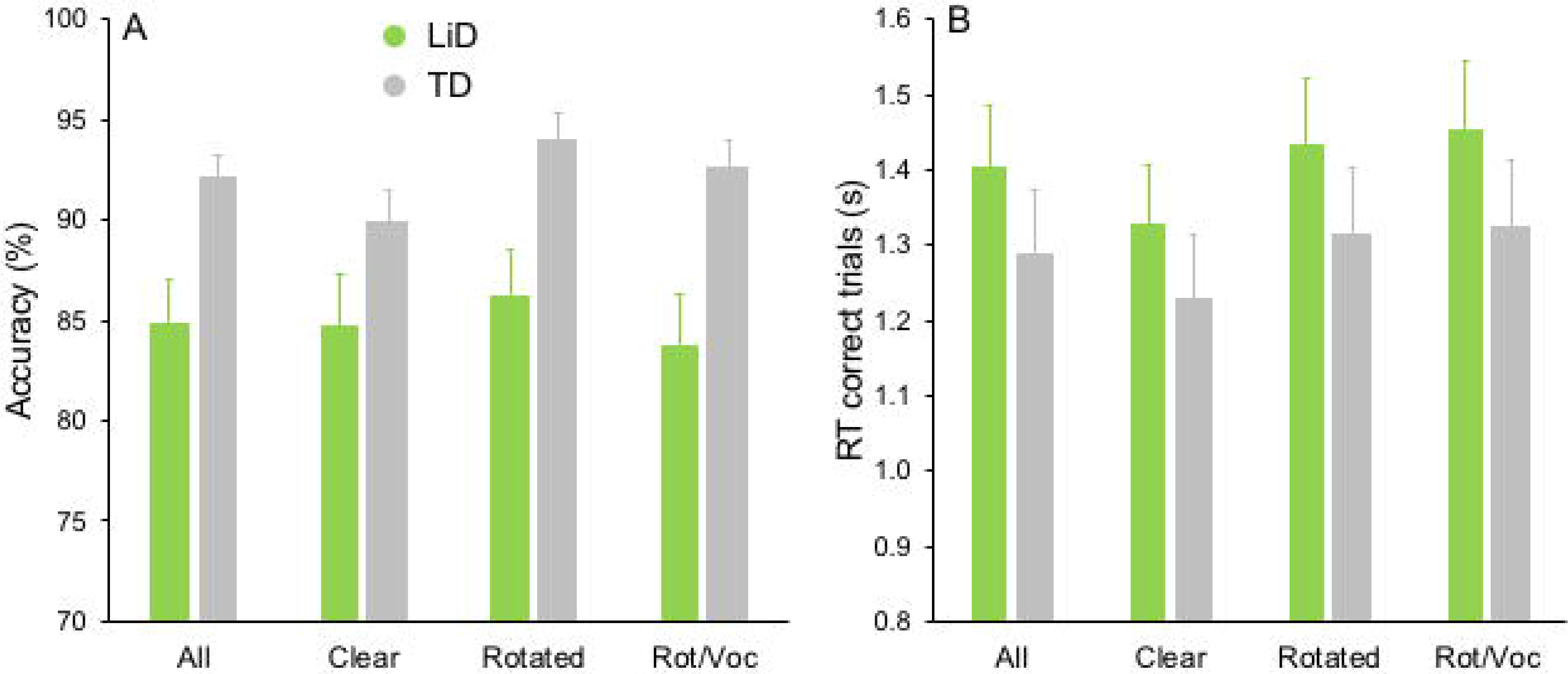
TD group showed superior performance on MRI real/unreal speech discrimination. **(A)** Mean (± sem) percent accuracy across sentences and (B) reaction times (RT, in seconds) for correct trials.

Neural activity associated with listening to the Speech, Phonetic and Intelligibility of the sentences did not significantly differ between the two groups. BOLD activation from across all children (regardless of group) are shown in Figure 8.

**FIGURE 8:**
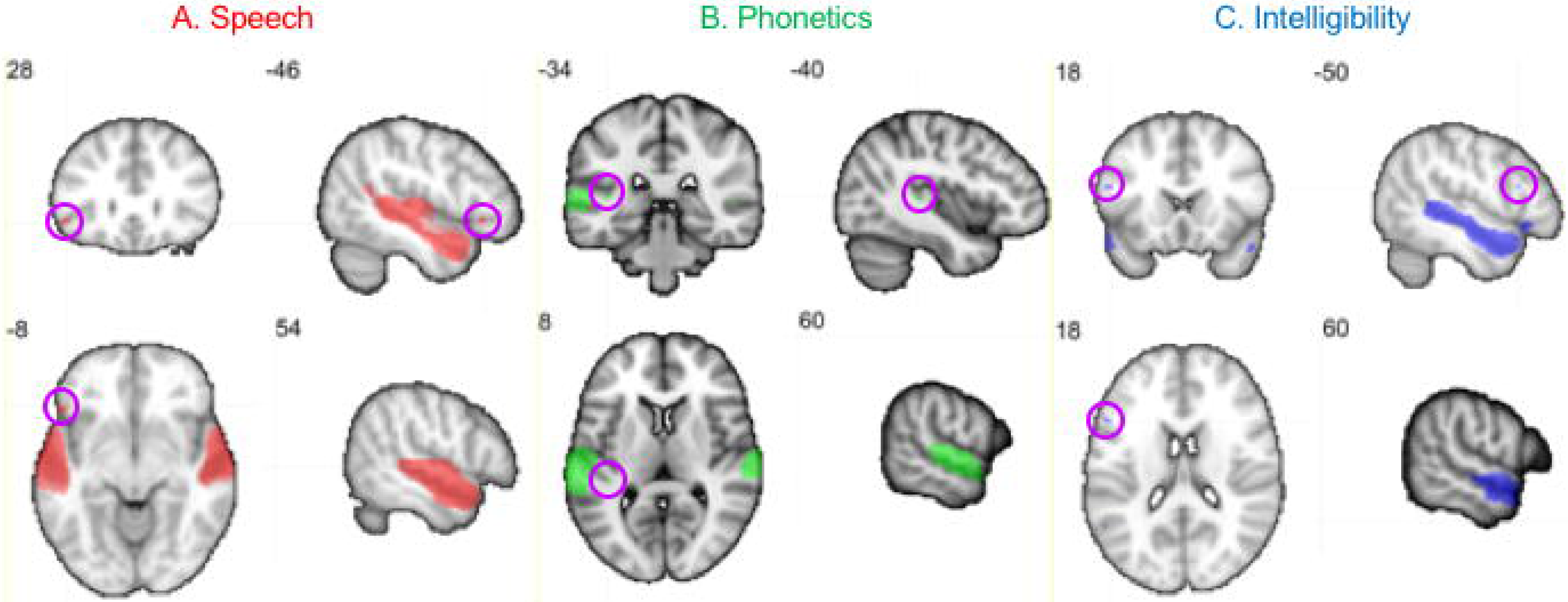
**Activation produced by Speech, Phonetic and Intelligibility processing during the speech categorization task**. Family-wise error correction of p < .001 and clustering threshold of k = 4 voxels was applied to create key ROIs for correlations with DL behavioral outcomes. Images are in neurological orientation and slice values refer to MNI coordinates. BrainNet software was used to display foci (Xia et al., 2013).

Activation patterns for Speech included bilateral auditory cortices (middle temporal gyrus, superior temporal gyrus and Heschl’s gyrus), planum temporale, left temporal fusiform cortex, inferior temporal gyrus, orbitofrontal cortex (OFC) and right parahippocampal gyrus. In the left OFC, a significant correlation was observed among the TD children between BOLD activation and the BDLT FR0 attention condition (Figure 9A). In contrast, children with LiD lacked such a correlation.

**FIGURE 9:**
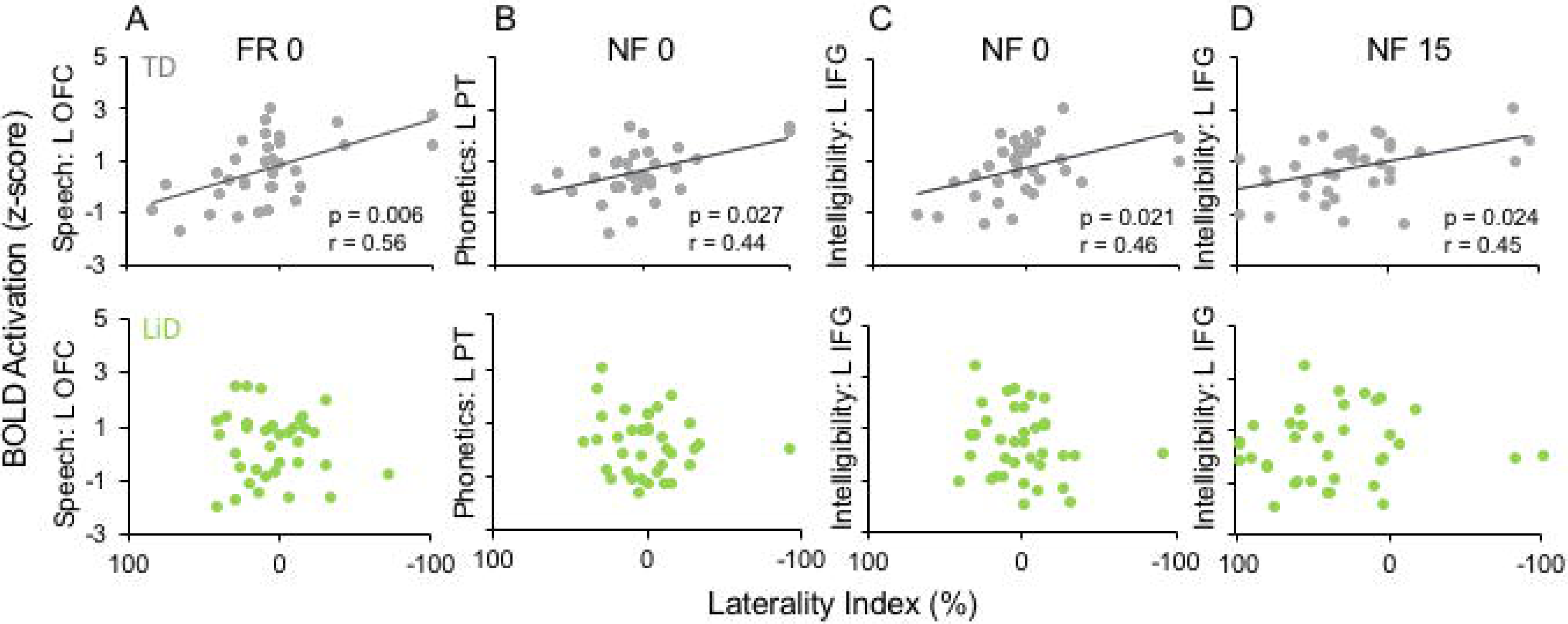
Selected, significant correlations between BDLT LI and brain activation in ROIs (Fig. 8), focusing on significant activation in TDs and equivalent ROI non-activations in LiD. All p-values in this figure have been adjusted to compensate for multiple comparisons. Further details in text.

Phonetic activation was seen in a more restricted region of posterior AC in the STG and planum temporale bilaterally and right Heschl’s gyrus (Figure 8B). Left planum temporale (PT) activation was correlated with the BDLT LI in TD children for the NF0 attention condition (Fig. 9B). For children with LiD, the relationship flipped, but not to the extent of a significant correlation.

The Intelligibility contrast revealed increased activation in the superior temporal gyrus, with a long anterior to posterior profile from the left temporal pole and along the STG and a more anterior temporal pole locus on the right (Fig. 8C). Significant correlations with BDLT LI were seen with the left IFG under both NF0 and NF15 conditions in the TD children. Again, no such correlations were observed in the children with LiD (Fig. 9C, D).

## DISCUSSION AND CONCLUSIONS

### Listening difficulties

Children with LiD performed normally in the BDLT Listen (non-forced) mode, as hypothesized, but they also performed normally in the BDLT Concentrate (FL, FR) modes, despite significantly impaired performance on speech-in-noise and cognitive tests. This is a new finding since most previous research on auditory processing differences between TD and non-TD children has focused on specific impairments in processing capacity of the left hemisphere, reflected in differential scores in the FL mode (Westerhausen & Hugdahl, 2010). The normal performance of children with LiD in the ILD 0 dB condition suggests that their cognitive insufficiency did not prevent them performing the DL task. Moreover, no significant differences were found between the groups on the right ear or left ear scores, suggesting no systematic hemispheric processing differences. Rather, the children with LiD were found to have a generalized disability to benefit from interaural level differences between the dichotic stimuli. The small REA found in both groups is consistent with weak REAs reported in previous studies of young children (Passow et al., 2013).

Performance on BDLT of children with LiD was more affected by varying ILD than was that of TD children. This could be because the children with LiD had a primary auditory problem, or that they were less able to offset greater sound level at either ear through attention modulation (Westerhausen & Hugdahl, 2010). Poor LiSN-S performance, particularly on the spatial advantage measure, may indicate a binaural interaction problem (Cameron & Dillon, 2008; Cameron et al, 2014; Glyde et al, 2013). Correlations between LI and LiSN-S advantage measures at high ILD support this interpretation, but LiD and TD groups did not differ in this respect. Inattention is a primary symptom of LiD, although its relationship to APD is controversial (Moore et al., 2010; Moore, 2018). However, there seems to be general agreement that many if not most children undergoing APD evaluation have attention difficulties that, at least, need to be taken into account by the examining audiologist (American Academy of Audiology, 2010).

### Age

There have been few studies of children using the BDLT. Age effects in this study generally mirrored those previously reported (Hugdahl et al., 2001; Passow et al., 2013), although the REA tended to get larger with increasing age, in contrast to the recent, NF-only study of (Kelley & Littenberg, 2019). In fact, comparison between current LiD data and adult data of (Westerhausen et al., 2009) suggested a robust, consistent increase in right ear influence with age, across ILD, supporting a “right ear weakness” hypothesis for LiD. This contrasted with TD children who had more of a “REA amplification” pattern of development where changes in LI with increasing age were asymmetric between left- and right-leading ILDs.

### fMRI

Both groups of children (LiD, TD) used the same brain areas to perform the sentence-listening task but relations between brain area activation and BDLT LI suggested the areas are used differently by the two groups. Left OFC was related to LI forced attention for TD but not for LiD. Other areas (left PT and IFG) also related to LI activation in TD, but not LiD. These results were all somewhat independent of ILD or task type (non-forced or forced). It was predicted that if the LiD children have an auditory processing deficit, we would find similar relationships between cortical activity and behavioral results for the BDLT attention conditions (i.e. FL and FR) and different relationships for the intensity manipulations in the BDLT (i.e. −15, 0 and 15). The reverse would suggest the LiD children have language processing deficits.

A lack of group differences in BOLD activations for the sentence-listening task in the MRI scanner suggests that the groups of children do not differ in the brain areas used to listen to language. However, the relationships between these brain areas and BDLT laterality suggest that these brain areas are used differently by each group. In the TD group, activation in a specific cortical area used for top-down processing of speech (left orbitofrontal cortex; OFC) was related to degree of laterality on a BDLT forced attention condition. However, this relationship was not found in the LiD group. Similarly, activation in areas used for bottom-up integration and synthesis of information underlying language (sound, phonetics and intonation in the left planum temporale and inferior frontal gyrus) was related to laterality of non-forced attention at different intensity levels in the TD group, but not in the LiD group. Lesser relationships were observed in other areas, with an overall pattern of some limited correlations with laterality among the TD group and lack of correlation in the LiD group.

Different group relationships were found between cortical activity and BDLT behavioral results for the attention conditions and also in the intensity manipulations. This suggests that the LiD children do not have clean cut auditory processing or language processing difficulties. These results do not indicate a redistribution of cortical listening areas in children with LiD but, instead, a reorganization as to how these areas are facilitated during language listening, with higher engagement of cortical listening areas associated with better listening task performance by the TD children.

### Implications for listening difficulties in children

Clinical use of dichotic assessment for APD in children has mostly used dichotic digits (Emanuel et al., 2011; Musiek, 1983). As discussed in the Introduction, the results of that assessment do not distinguish between an auditory and a cognitive explanation of those children’s LiD, although the test may correctly identify children without hearing loss as having an auditory perceptual or speech coding problem. In other studies, using the less cognitively but more auditorily demanding BDLT, older children and adults with a wide variety of learning, neurological and mental health diagnoses had a generally weak left ear performance in the FL condition (Westerhausen & Hugdahl, 2010). This was interpreted as a means for testing top-down executive function that we found here in the Toolbox data to be impaired in children with LiD. However, we did not find a consistent poor performance on the FL task in that group.

A number of observations have been made about dichotic ear advantages in children with APD (Moncrieff et al., 2017). Some studies have focused on the prevalence of a REA or LEA, suggesting balance between the two is initially more even but unstable, but that a consistent REA emerges with age (Moncrieff, 2011). However, the current results, and others (Hugdahl et al., 2001) show that the absolute level of LI increases (i.e. larger LEA and REA) with age, contrary to the report of (Moncrieff, 2011), and that use of a binary LEA/REA distinction can be misleading. Other results in adults have shown that larger LIs, either positive or negative, are associated with better accuracy on the BLDT (Hirnstein et al., 2014).

A new term, “amblyaudia” was introduced by (Moncrieff et al., 2016) to designate “an abnormally large asymmetry between the two ears during dichotic listening tasks with either normal or below normal performance in the dominant ear”. The results of the study reported here only partially supported amblyaudia in the children with LiD. Their performance was statistically indistinguishable from that of the TD children at ILD 0 dB, the usual condition for testing. But, consistent with the definition, there was a larger than normal asymmetry in the ILD 15 dB conditions, with normal or below normal number of correctly reported digits in the right ear and an above normal number of correct reports in the left ear. In a review, Whitton & Polley (2011) discussed amblyaudia in the context of long-term effects of conductive hearing loss on auditory system plasticity, induced in children predominantly by otitis media. While building a convincing case from the literature for such plasticity, the relevance of such findings to the children in this study is unclear; a similar proportion of children in each group had a history of PE tubes (Hunter et al., 2019).

Several forms of dichotic training have been proposed, and we know of at least two that are in current evaluation or practice for the treatment of amblyaudia (DIID - Musiek, 2004); ARIA - Moncrieff et al., 2017) and other abnormalities detected through dichotic listening evaluation (Emanuel et al., 2011). Unfortunately, it remains unclear what sort of benefits might be obtained from such training or whether any of the proposed methods generalize to improved listening in real-world challenging environments. (Hugdahl et al., 2009) present several arguments *against* training using the BDLT to treat impaired performance on the FL instruction task. These arguments are, briefly, that the DL task is very simple and therefore unchallenging, that it shows little or no learning effect, and that executive functions are not amenable to training. It is unclear what form of training might be useful for normalizing specific DL behavior patterns of the children with LiD in the current study. However, interventions that improve auditory attention should be generally useful for these children.

## ACKNOWLEDGMENTS

We wish to thank all the families who are participating in the SICLID study at the Listening Lab of Cincinnati Children’s Hospital Medical Center. This research was supported by NIH grant R01DC014078 and by the Cincinnati Children’s Research Foundation. David Moore is supported by the NIHR Manchester Biomedical Research Centre.

